# Temporal seascape genomics identifies evolutionarily significant units in a highly exploited marine resource, the wedge clam *Donax trunculus*

**DOI:** 10.64898/2025.12.03.691988

**Authors:** Benestan Laura, Marc Baeta, Carlos Saavedra, Marina Delgado, Ana Insua, Silvia Falco, Miguel Rodilla, Luis Silva, Miriam Hampel, Ciro Rico

## Abstract

Clam fisheries along the Spanish coast have declined unevenly, while stock management often follows political rather than biological boundaries. To inform conservation and fishery management, this study used temporal seascape genomics to assess population structure, genetic diversity and effective population size (*Ne*) in the wedge clam *Donax trunculus* across approximately 2,000 km of the Iberian coast. Using 7,830 putatively neutral and 649 candidate outlier SNPs from 331 individuals, the analyses confirmed broad Atlantic–Mediterranean differentiation, identified the Alboran Sea as a distinct genetic unit, and revealed finer-scale Atlantic substructure. Temporal genomic comparisons showed that population structure remained largely stable over the study period, although genetic diversity changed in a site-specific manner across fishing areas. Genotype–environment analyses indicated that spatial structure explained more genomic variation than environmental predictors, especially after accounting for spatial– environmental collinearity. Overall, the results show that temporal seascape genomics can distinguish persistent biogeographic structure from short-term, local genomic change in exploited marine invertebrates, providing a robust framework for biologically informed stock delineation. The genetically distinct groups identified here should therefore be considered separately in conservation, fishery management and any future reinforcement planning.

## Introduction

The overexploitation of commercially valuable marine species, together with climate change and other anthropogenic stressors, is a major driver of biodiversity loss and can reduce the demographic and evolutionary resilience of exploited populations (Hald-Mortensen, 2023; Jaureguiberry *et al*, 2022; Thomson *et al*, 2021). Bivalves such as clams, cockles and scallops are particularly vulnerable because many support intensive small-scale fisheries while also providing ecosystem services and socioeconomic value to coastal communities (Azmi *et al*, 2022; McLaverty *et al*, 2020). Population declines can reduce census size, alter recruitment dynamics and, when sustained over sufficient generations, erode genetic diversity and effective population size (*Ne*), thereby limiting adaptive potential under continued environmental change (Petit-Marty *et al*, 2022; Sadler *et al*, 2023). However, the genomic consequences of exploitation are not always immediate or uniform, especially in marine invertebrates with high fecundity, planktonic larval dispersal and potentially extensive gene flow. Temporal genomic approaches are therefore valuable because they allow direct assessment of whether allele frequencies, genetic diversity and population structure have changed across management-relevant timescales (Clark *et al*, 2023; Snead and Clark, 2022).

Accurate delineation of management units is central to sustainable fisheries management. In exploited marine resources, management boundaries often follow administrative jurisdictions rather than biological population structure, which can lead either to overharvesting of discrete stocks or to unnecessary subdivision of genetically connected populations (Benestan, 2020; Cadrin *et al*, 2010; Punt, 2023). Genome-wide markers can improve the delineation of biologically meaningful management units by resolving both neutral population structure, which primarily reflects demographic history, drift and gene flow, and putatively adaptive variation, which may reveal environmentally associated differentiation relevant to conservation and any future population-reinforcement planning (Gagnaire *et al*, 2015; Liggins *et al*, 2019). Seascape genomics provides an explicit framework for this purpose by integrating genomic, geographic and environmental information to assess how migration, genetic drift, biogeographic barriers and spatially structured environmental gradients jointly shape genetic diversity across marine landscapes (Liggins *et al*, 2019; Riginos and Liggins, 2013). However, because environmental gradients often co-vary with geographic distance and oceanographic discontinuities, seascape-genomic interpretations require caution and are strongest when neutral and outlier-based patterns are evaluated together. In this context, the wedge clam *Donax trunculus* provides a valuable system for testing whether broad biogeographic units previously inferred from microsatellites are temporally stable, whether genome-wide neutral and putatively adaptive SNPs refine those units, and whether site-specific temporal changes in genetic diversity and effective population size are associated with contrasting local histories of fishery closure, reduced harvesting pressure and continued exploitation.

The wedge clam is a commercially and culturally important bivalve distributed along the eastern Atlantic coast from Senegal to France and throughout the Mediterranean and Black Sea (Baeta *et al*, 2023; Delgado and Silva, 2018; Ramón, 1993). Across the Iberian Peninsula, it supports small-scale fisheries that have experienced uneven trajectories over recent decades. The Spanish Mediterranean has undergone a sharp decline in official landings, from 1,181.22 tonnes in 1994 to 37.04 tonnes in 2024 (Baeta *et al*, 2025), leading to formal closures or strong restrictions in several areas, including the Gulf of Valencia and northern Catalonia (Escrivá *et al*, 2021). In other areas, including central Catalonia and the Almería coast, fishing activity has been progressively abandoned because low stock density has reduced profitability. Almería is referred to here as a fishery region; the Alboran sampling locality analysed genetically was Málaga. Galicia has also shown a persistent decline in reported landings since the early 2000s, whereas parts of the Andalusian Atlantic coast and Portuguese waters show more stable recent catches following earlier declines. Because landings can reflect a combination of abundance, regulation, effort and profitability, they should be interpreted cautiously as indicators of demographic change. These contrasting regional trajectories make *D. trunculus* an informative system for assessing whether fishery closures and reduced harvesting pressure are accompanied by detectable genomic changes.

Previous genetic studies using microsatellites and mitochondrial DNA identified a major Atlantic-Mediterranean break and supported the recognition of broad management units corresponding to the Atlantic Ocean, Alboran Sea and north-western Mediterranean/Balearic Sea (Fernández-Pérez *et al*, 2018; Marie *et al*, 2016; Nantón *et al*, 2017). These studies also reported relatively weak differentiation along the Atlantic Iberian coast, suggesting substantial connectivity among Atlantic localities. However, previous analyses were based on comparatively low-density marker sets and did not evaluate temporal change using genome-wide SNP data. In addition, the extent to which adaptive or environmentally associated variation refines these broad units remains unresolved.

Here, we used temporal seascape genomics to address five linked hypotheses about the spatial and temporal genomic structure of *D. trunculus* across the Iberian Peninsula. First, we tested whether genome-wide neutral SNPs recover the broad management units previously inferred from microsatellites, corresponding to the Atlantic Ocean, Alboran Sea and north-western Mediterranean/Balearic Sea. Second, we tested whether neutral and candidate outlier SNPs provide additional resolution within these broad units, particularly along the Iberian Atlantic coast. Third, we assessed whether population structure remained temporally stable between pre- and post-closure samples, as expected if high fecundity, planktonic larval dispersal and gene flow buffer short-term allele-frequency change. Fourth, we examined whether genetic diversity and effective population size show region- or site-specific temporal shifts associated with contrasting histories of closure, reduced fishing pressure and local recruitment dynamics. Finally, we tested whether spatial structure explains genomic variation more strongly than environmental predictors once spatial-environmental collinearity is considered, and whether candidate environmentally associated variation can inform spatial management and future reinforcement planning.

## Methods

### Sampling design and filtering steps

The sampling design covers almost the entire Spanish range of *D. trunculus*. Sample collection procedures followed those described in Marie *et al* (2016) and Nantón *et al* (2017). Samples were collected from seven locations spanning the Atlantic and Mediterranean coasts of Spain, with one additional site in Portugal (Setúbal) (Table 1, Figure 1). Temporal replicates were obtained in 2020, 2022 and 2023 from the same geographic regions sampled by Marie *et al* (2016) and Nantón *et al* (2017) although logistical constraints prevented resampling at the exact same coordinates in all cases. In such instances, samples were collected within approximately 10 km of the original sampling sites. We consider this spatial offset unlikely to affect the interpretation of temporal genetic patterns for two reasons. First, both previous studies reported no detectable genetic differentiation among nearby localities within the same regional genetic units, indicating genetic homogeneity at this spatial scale. Second, given the planktonic larval phase of *D. trunculus* which is approximately 21 days according to Ruiz-Azcona *et al* (1996), distances of a few kilometres are expected to be negligible relative to the scale over which gene flow occurs. Therefore, while the temporal comparisons should be interpreted as regional rather than exact site-level resampling, the small spatial displacement among sampling points is unlikely to generate artefactual differences in genetic structure or diversity. A total of 331 *D. trunculus* individuals were genotyped using the DArTseq™ platform. To ensure high-quality SNP data, we applied a series of filtering steps based on quality metrics provided by the DArTseq™ pipeline, implemented in R using the *dartR* (Gruber *et al*, 2018) and *poppr* (Kamvar *et al*, 2014) packages. Loci were retained based on the following criteria: a minimum allele frequency (MAF > 0.05), a maximum missing data rate of 35% per locus and per individual, and a pairwise linkage disequilibrium threshold of R² < 0.8 (see Table S1). Relatedness estimates were calculated to identify and remove full siblings (r2 > 0.4) or half-siblings (r2 > 0.2) using the R package *related* (Pew *et al*, 2015).

**Figure 1.**
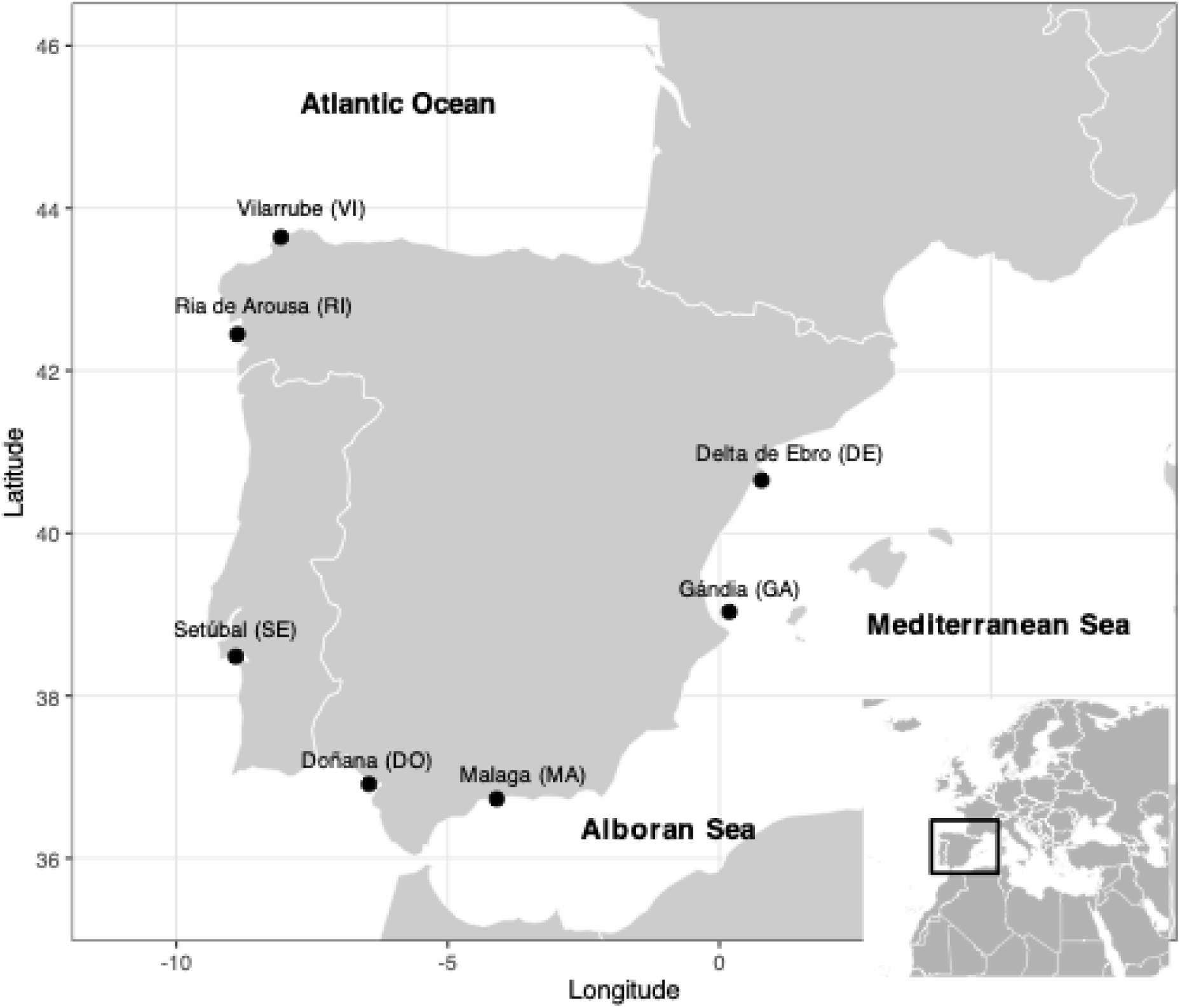
Sampling map. Map showing the distribution of the seven locations sampled.

**Table 1.** Sampling locations with their corresponding site code, year of sampling, geographic coordinates and number of samples successfully genotyped.

| Sampling locations | Code | Latitude | Longitude | Year of sampling | n |
| --- | --- | --- | --- | --- | --- |
| Vilarrube | VI | 43.64250000 | -8.07500000 | 2011 | 29 |
|  |  |  |  | 2020 | 27 |
| Ría de Arousa | RI | 42.44750000 | -8.87555556 | 2014 | 25 |
|  |  |  |  | 2023 | 31 |
| Setúbal | SE | 38.48222222 | -8.90416667 | 2014 | 28 |
| Doñana | DO | 36.91222222 | -6.45166667 | 2014 | 26 |
|  |  |  |  | 2022 | 30 |
| Málaga | MA | 36.73027778 | -4.09944444 | 2023 | 28 |
| Gandía | GA | 39.03361111 | -0.18305556 | 2014 | 21 |
|  |  |  |  | 2022 | 26 |
| Delta del Ebro | DE | 40.65027778 | 0.77888889 | 2014 | 31 |
|  |  |  |  | 2022 | 29 |

A putatively neutral dataset was subsequently generated by removing loci that deviated significantly from Hardy-Weinberg equilibrium (HWE) in more than 70% of the sampling locations, using the *gl.filter.hwe* function in the *dartR* package. Importantly, this step was conducted after outlier locus detection to avoid inadvertently discarding loci potentially under selection, which frequently deviate from HWE expectations. This filtering produced three analytical datasets: all retained SNPs after quality control, a putatively neutral dataset after removal of candidate outlier loci, and a candidate outlier dataset. These datasets showed high reproducibility and were used for downstream analyses as appropriate.

### Outlier detection

Outlier loci putatively associated with spatially heterogeneous selection were identified using two complementary approaches. The first method employed pcadapt (Luu *et al*, 2017), which detects loci potentially under selection by analysing genotype data at the individual level. Briefly, this method performs a principal component analysis (PCA) to capture the major axes of genetic variation and then identifies SNPs that are strongly correlated with these axes, suggesting possible adaptive divergence. For each SNP, a q-value was computed as a multiple-testing-corrected p-value, thereby controlling the false discovery rate associated with the large number of SNPs tested. SNPs with a q-value < 0.05 were classified as candidate outliers.

The second approach used OutFLANK, an R package implementing the method developed by Whitlock and Lotterhos (2015). OutFLANK detects loci under selection by estimating the distribution of *F_ST_* expected under neutrality. The method fits a likelihood model to a trimmed distribution of observed *F_ST_* values, deliberately excluding extreme values that may represent loci under selection. From this inferred neutral distribution, q-values were calculated for each locus. After multiple-testing correction, pcadapt identified 649 candidate loci, whereas OutFLANK did not detect significant outliers. Consequently, downstream outlier analyses were based on the 649 pcadapt candidate loci and are interpreted cautiously as putatively adaptive candidates rather than as high-confidence loci independently supported by both methods.

### Population structure at both neutral and outlier loci

Population structure was assessed separately for the neutral and candidate outlier SNP datasets using two complementary clustering approaches. First, we applied a non-model-based *K*-means clustering algorithm using the *find.cluster* function from the *adegenet* package (Jombart *et al*, 2008). This method estimates the optimal number of genetic clusters (*K*) by assessing the Bayesian Information Criterion (BIC) across a range of increasing K values. Prior to clustering, genotype data were transformed using Principal Component Analysis (PCA), and clustering was performed without assuming any underlying population genetic model.

Second, we conducted a model-based clustering analysis using sparse non-negative matrix factorisation (sNMF) implemented in the *LEA* R package (Frichot and François, 2015). This method, analogous to STRUCTURE (Pritchard *et al*, 2000), estimates individual ancestry coefficients and infers population structure assuming Hardy-Weinberg equilibrium. The optimal number of clusters was determined using the cross-entropy criterion. Briefly, this criterion operates randomly masking a proportion of genotypes (typically 10%) in the dataset, fitting the sNMF model on the remaining data, and evaluating how accurately the masked genotypes can be predicted from the inferred ancestry coefficients and allele frequency estimates. This procedure is repeated across a range of *K* values, and the value of *K* minimising the cross-entropy - reflecting the best predictive accuracy - is retained as the optimal number of clusters. Ten independent runs were performed for each value of *K* to account for stochastic variation in the optimisation algorithm, and the run with the lowest cross-entropy was retained for further analysis.

Based on the population grouping identified through using clustering analyses, we computed analyses of molecular variance (AMOVA) using *poppr* package in R (Kamvar *et al*, 2014). The significance of variance components was tested with 20,000 permutations. Populations were grouped into hierarchical sets reflecting the clustering results.

### Temporal population structure

Temporal changes in genetic structure and diversity were assessed using samples collected over multiple time points at two Mediterranean locations - Delta del Ebro (DE) and Gandía (GA) - and three Atlantic locations - Doñana (DO), Ría de Arousa (RI) and Vilarrube (VI). These sites were selected to evaluate the genetic consequences of fishery closures, reduced harvesting pressure and localised cessation of fishing activity in Spain (see Table 2). To assess temporal stability of population structure, we conducted a supervised Discriminant Analysis of Principal Components (DAPC) using the *dapc* function in the *adegenet* R package (Jombart *et al*, 2010). Sampling location was used as the prior grouping variable (grp). The number of principal components to retain was determined through the α-score optimization, which indicated that the first 15 PCs and nine discriminant factors were retained in the analysis.

**Table 2.** Analysis of Molecular Variance (AMOVA)

| Source of variation | %var | F-stat | df | P-value |
| --- | --- | --- | --- | --- |
| Within samples | 75.11 | F<sub>IS</sub> | 331 | - |
| Between samples within population | 19.16 | F<sub>ST</sub> | 319 | 0.248 |
| Among populations within groups | 0.5 | F<sub>SC</sub> | 10 | 0.005 |
| Among groups | 5.22 | F<sub>CT</sub> | 1 | 0.001 |

### Temporal change in genetic diversity

Genetic diversity metrics were calculated for each sampling location and time point using the *hierfstat* package, including observed heterozygosity (*Ho*), expected heterozygosity (*He*) and allelic rarefaction (AR). Pairwise genetic differentiation among all sampling locations and time points was quantified using *F_ST_* values based on Weir and Cockerham (1984) estimator, computed with 10,000 bootstrap replicates and bias-corrected 95% confidence intervals, as implemented in the *dartR* package.

### Estimating change in effective size

Contemporary effective population sizes (*Ne*) were estimated for each genetic cluster and sampling year using the linkage disequilibrium method implemented in NeEstimator v2 (Do *et al*, 2014), with a MAF threshold of 0.05, and all other parameters set to default. Confidence intervals were calculated using the biased-corrected parametric method, following Waples (2006). Briefly, this approach corrects for the upward bias inherent in *Ne* estimation from finite samples, which arises because the observed mean *r²* (a measure of linkage disequilibrium between pairs of loci) is inflated relative to its expectation when sample sizes are small. The bias correction adjusts the expected *r²* by accounting for the sampling variance introduced by a finite number of individuals and loci. Parametric confidence intervals are then derived from the chi-squared distribution of the corrected *r²* statistic, assuming normality of the estimator. This method is preferred over jackknife approaches when sample sizes are moderate, as it provides more accurate interval coverage (Waples, 2006; Waples and Do, 2010).

We also applied the temporal method to estimate Ne for locations with temporal replicates for which robust estimates were retained: Delta del Ebro (DE), Doñana (DO), Gandía (GA), Ría de Arousa (RI) and Vilarrube (VI). This approach estimates Ne from changes in allele frequencies observed between two sampling points. Given that generation time in D. trunculus has been reported to range from one to two years, the two sampling intervals of approximately 5 to 10 years correspond roughly to 4 to 10 generations, depending on local conditions and the generation time assumed. Three estimators were implemented in NeEstimator v2 (Jorde and Ryman, 1995; Nei and Tajima, 1981; Pollak, 1983). A MAF threshold of 0.05 was applied consistently across all analyses to reduce random noise and computational load.

### Spatial variables

To investigate isolation by distance (IBD), we tested the null hypothesis of no correlation between pairwise genetic differentiation [*F_ST_* /(1- *F_ST_*)] and in-water geographic distance (*i.e.* within coastline distances). IBD was tested using *mantel.rtest* function from the *ape* package, with 1,000 random permutations (Paradis *et al*, 2004). In-water geographic distances were calculated using bathymetry maps imported with the *get.NOAA* function from the *marmap* R package. Least-cost marine path (i.e., shortest paths in water, avoiding land) were computed using the *trans.mat* and *lc.dist* functions from the same package.

To further characterise spatial structure, we performed a distance-based Moran’s eigenvector maps (dbMEM) analysis using the *dbmem* function from the *adespatial* R package (Dray *et al*, 2019). dbMEMs represent a spectral decomposition of the spatial relationships among sampling locations, generating (n - 1) orthogonal eigenfunctions that capture spatial patterns at multiple scales (Dray *et al*, 2006). dbMEM variables were derived from the in-water distance matrix. We additionally applied the PCNM function from the *PCNM* R package, using 1,000 permutations, which transforms geographic distances into a set of orthogonal spatial variables suitable for use in constrained ordination analyses (Legendre and Gallagher, 2001) enabling the explicit incorporation of spatial structure into downstream statistical models.

### Environmental variables

Environmental variables - including sea surface temperature (SST), sea surface salinity (SSS), sea floor temperature (SFT), dissolved inorganic carbon, chlorophyll, nitrate, phosphate, dissolved oxygen, iron, pH, phytoplankton - were retrieved from the Marine Copernicus database (https://marine.copernicus.eu/). For each variable, monthly minimum, maximum and mean values were extracted for the period 2010-2023. This timeframe encompasses environmental variation over at least six to twelve generations, based on the generation time of *D. trunculus*, which typically is about one-two years, depending on environmental conditions such as temperature, salinity, and primary productivity, as well as geographic location (Ansell and Lagardère, 1980; Guillou, 1980; Lamine *et al*, 2020; Mazé and Laborda, 1988; Ramón, 1993; Ramón *et al*, 1995; Zeichen *et al*, 2002).

To incorporate proxies of anthropogenic pressure, we calculated human population density (PDNE) within a 25 km radius surrounding each sampling location. This was derived using shapefiles downloaded from the Global Multi-Resolution Topography (https://www.gmrt.org/GMRTMapTool/) and population data from the NASA earth observations (https://neo.gsfc.nasa.gov/). Additionally, we estimated the percentage of surrounding land surface (LAND) within the same 25 km radius around each site to account for coastal topography, which may influence larval retention. Sites with higher percentage of surrounding land were interpreted as more enclosed systems, potentially exhibiting greater larval retention compared to more open coastal sites.

### Redundancy Analysis

Redundancy analysis (RDA) is a multivariate ordination technique that extends linear regression to multivariate response data - in this case, individual genotypes across multiple SNPs. First, we calculated individual Euclidean genomic distances using the *dist* function available in the R package *adegenet* (Jombart *et al*, 2008) and a principal coordinate analysis (PCoA) was subsequently performed on the resulting distance matrix. The resulting principal component (PC) axes, representing the position of individuals in the genomic multivariate space, were used as response variable in a distance-based RDA (hereafter dbRDA). Explanatory variables included spatial structure, represented by Moran’s eigenvector maps (MEMs) and the set of environmental variables described above (Borcard and Legendre, 2002). Prior to model fitting, collinearity among explanatory variables was assessed, and a final parsimonious model was defined by removing non-significant and highly collinear variables through forward selection. All code used for the dbRDA and dbMEM analyses is available at https://github.com/laurabenestan/db-RDA-and-db-MEM.

## Results

### SNP filtering

The initial unfiltered SNP dataset generated by DArT comprised 378 individuals and 125,301 loci. Following quality control filtering, 47 individuals were removed due to low call rate (<65%) and 116,822 loci were discarded based on the following criteria: minor allele frequency (MAF < 0.05), excess heterozygosity (H_O_ > 0.5), and missing data exceeding 65%. The number of SNPs at each step is reported in Table S1. The final overall neutral dataset retained 331 individuals and 7,830 SNPs. A second dataset restricted to the five temporal sites - Gandía (GA), Delta del Ebro (DE), Doñana (DO), Vilarrube (VI), and Ría de Arousa (RI) - included 7,805 putatively neutral SNPs genotyped in 205 individuals. Outlier detection analyses identified 649 candidate loci with pcadapt (Figure S1), whereas OutFLANK detected none after multiple-testing correction. Subsequent outlier analyses therefore used the 649 pcadapt candidate loci and should be interpreted cautiously.

### Broad scale neutral population structure

Using a dataset of 7,830 putatively neutral SNPs, K-means clustering identified two major spatial genetic clusters along the Spanish coastline, corresponding to the Atlantic region - Doñana (DO), Málaga (MA), Ría de Arousa (RI), Vilarrube (VI) and Setúbal (SE) - and the Mediterranean region - Gandía (GA) and Delta del Ebro (DE). In contrast, sNMF analysis detected three genetic clusters, further distinguishing the Alboran Sea samples - Málaga (MA) - as a separate genetic unit (Figure 2). The Analysis of Molecular Variance (AMOVA) based on the two genetic clusters identified by K-means revealed that most genetic variation was distributed within samples (75.11%). Variation among populations within groups was low but significant (0.50%, P = 0.005), whereas variation among groups accounted for 5.22% of the variance and was statistically significant at the conventional 0.05 threshold (P = 0.001; Table 2). Variation between samples within populations was comparatively high but non-significant (19.16%, P = 0.248).

**Figure 2.**
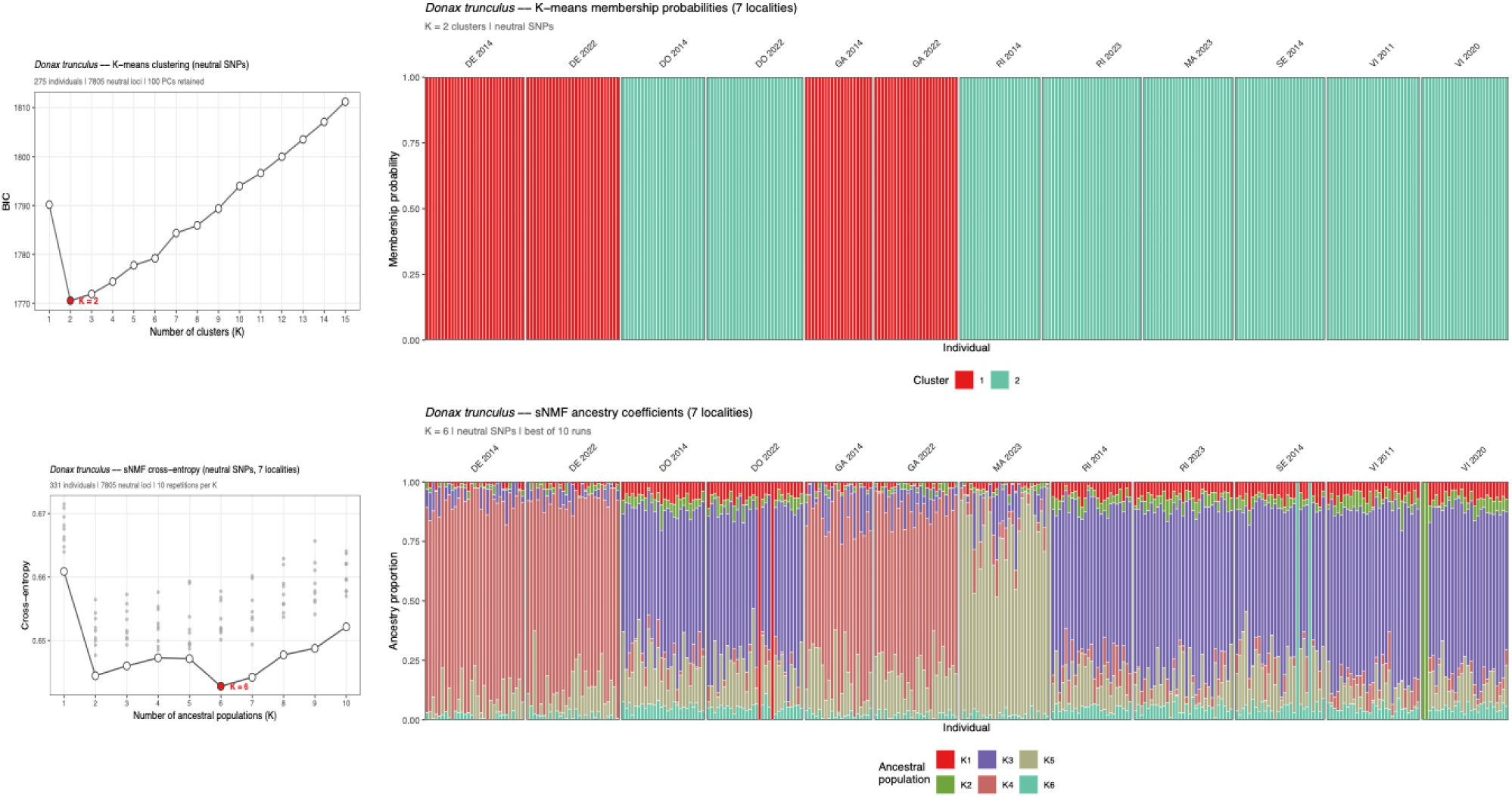
Broad-scale neutral population genomic structure inferred from 7,830 putatively neutral SNPs genotyped in 331 individuals. Bayesian Information Criterion (BIC) for K-means clustering and cross-entropy for sNMF are shown across values of K. Individual assignment probabilities are shown for the optimal clustering solutions, with colours indicating inferred genetic clusters.

### Broad scale potentially adaptive population structure

Analysis of the 649 pcadapt candidate outlier SNPs suggested additional spatial structure, but these results should be interpreted cautiously because OutFLANK did not detect significant outliers after multiple-testing correction. K-means clustering indicated six groups, including differentiation of the Alboran Sea and a subdivision within the Atlantic region between the southern Atlantic population (Doñana, DO) and central/northern Atlantic populations (Vilarrube, VI; Ría de Arousa, RI; and Setúbal, SE; Figure S1). The sNMF analysis also suggested fine-scale structure, although the number of inferred clusters differed from K-means and should therefore be treated as exploratory rather than as a definitive set of adaptive units (Figure S1). Individuals from Doñana (DO) showed partial assignment to both southern and central Atlantic components, consistent with its intermediate position within the Atlantic structure.

### Temporal population structure

This pattern was further corroborated by Discriminant Analysis of Principal Components (DAPC) conducted on the five sites with temporal replicates. The first discriminant axis, accounting for 82.2% of the between-group variance, clearly separated individuals from the Mediterranean Sea (GA, DE) from those originating from the Atlantic Ocean (DO, RI, VI), confirming the primary marine basin as the dominant axis of genetic differentiation (Figure 3). Notably, Mediterranean individuals formed a more compact cluster in DAPC space compared to their Atlantic counterparts, suggesting lower within-region genetic variability in the Mediterranean Sea. The second discriminant axis (15.5%) provided finer resolution within the Atlantic group, separating the southernmost population (DO), located along the southern Spanish Atlantic coast, from the central Atlantic populations (VI, RI), consistent with a pattern of latitudinal genetic structuring. The third discriminant axis explained little additional variance and did not reveal meaningful substructure among sampling sites.

**Figure 3.**
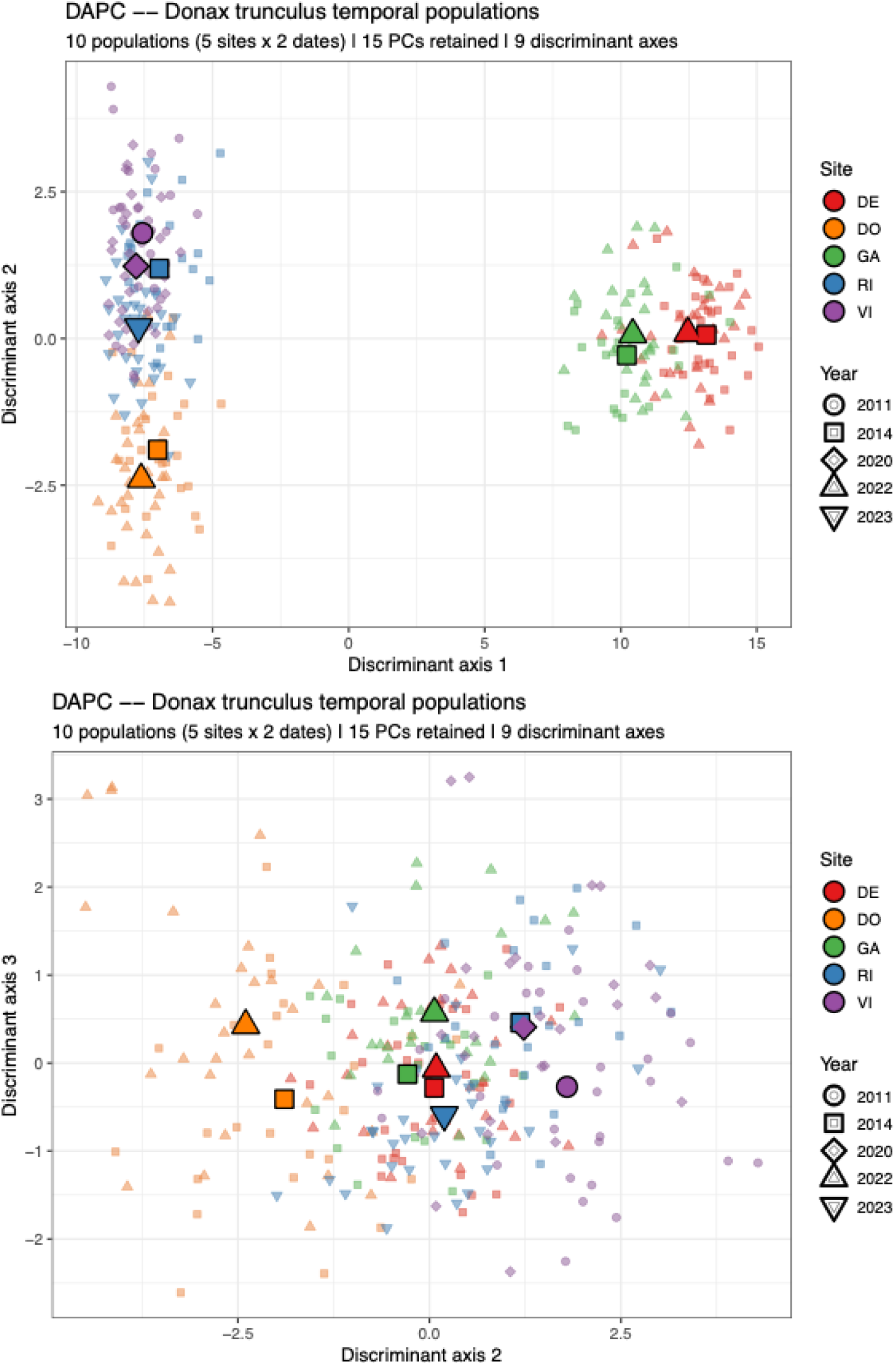
Temporal neutral population structure and pairwise differentiation. Discriminant Analysis of Principal Components was performed on 7,805 putatively neutral SNPs genotyped in 205 individuals sampled across sites and years. Each point represents one individual, colour-coded by sampling location and shape-coded by sampling year. Temporal overlap within locations indicates stability of population structure through time.

### Genetic differentiation

The temporal stability of the genetic structure was further supported by the DAPC results, in which temporal replicates from the same location consistently clustered together (Figure 3). The lowest pairwise *F_ST_* values were observed between temporal replicates, and most temporal comparisons were non-significant or only marginally significant under the selected threshold: RI_2014/RI_2023 (*F_ST_* = 0.0016, P-value = 0.01), VI_2011/VI_2020 (*F_ST_* = 0.0011, P-value = 0.06), DE_2014/DE_2022 (*F_ST_* = 0.0014, P-value = 0.02), and DO_2014/DO_2022 (*F_ST_* = 0.0010, P-value = 0.05). The negative *F_ST_* estimate for GA_2014/GA_2022 (*F_ST_* = -0.0005, P-value = 0.71) is interpreted as indistinguishable from zero, indicating no detectable temporal differentiation at this site. The highest and most significant levels of genetic differentiation were consistently detected between Atlantic and Mediterranean sites, reflecting limited gene flow across these two basins. At a finer spatial scale, pairwise comparisons among sites within each basin were statistically significant but generally low, suggesting restricted but detectable gene flow among locations. Isolation by distance (IBD) analysis across Atlantic sampling sites revealed a positive but non-significant correlation between genetic and geographic distances (R² = 0.593, P-value = 0.073; Figure 4) and should therefore be interpreted cautiously.

**Figure 4.**
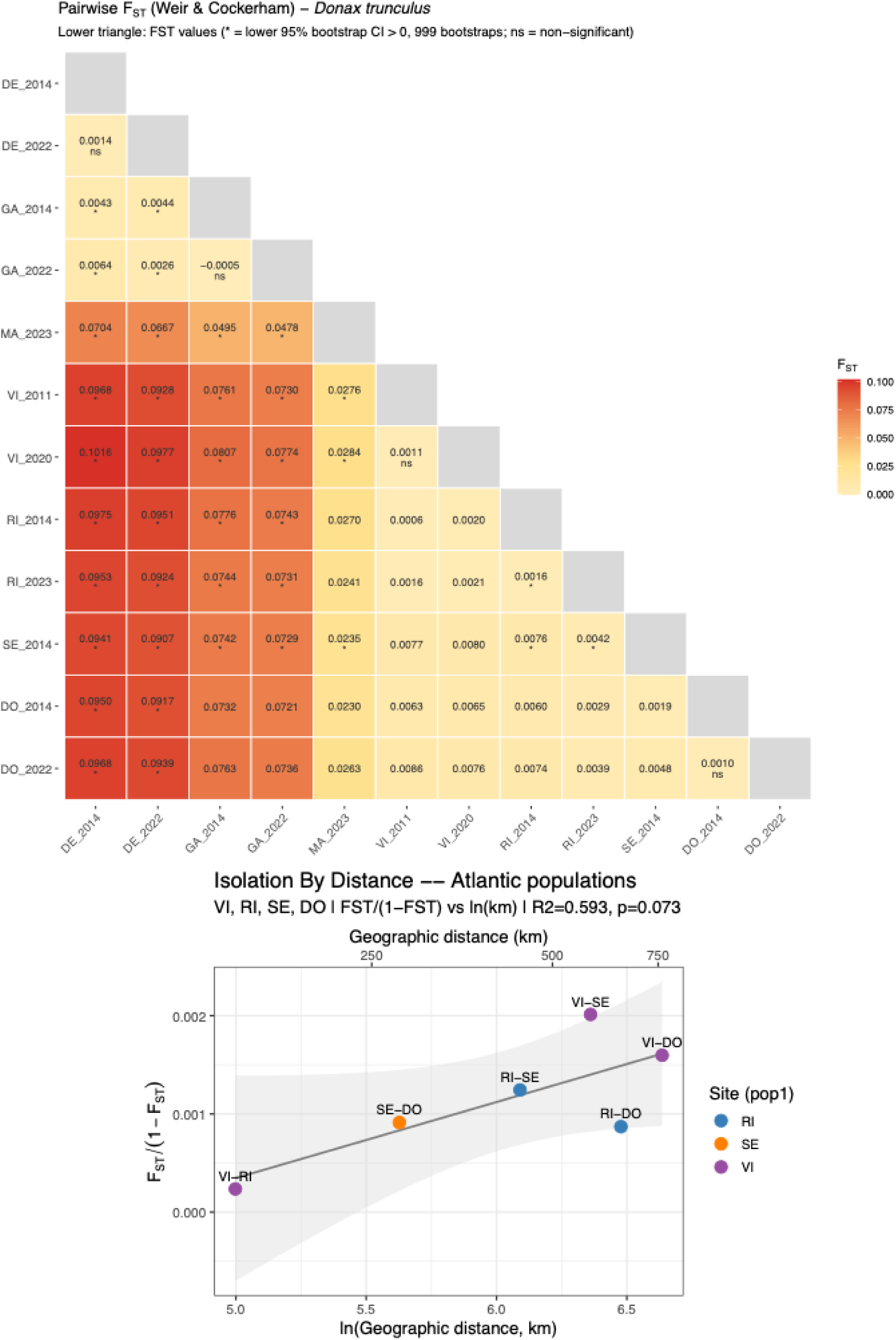
Genetic differentiation. (Top) Weir and Cockerham’s *F_ST_* index of genetic differentiation among pairwise sampling locations, coloured on a gradient from yellow (low values) to red (high values). Significant values are indicated with an asterisk. (Bottom) Exploratory isolation-by-distance analysis restricted to the four Atlantic sampling locations.

### Temporal stability of population structure

Overall, population structure remained temporally stable across all locations, as evidenced by a strong correlation between pairwise *F_ST_* values estimated before and after the fishery closure (R² = 0.8323; P-value < 0.001). The sole exception was the VI/RI comparison, which shifted from non-significant differentiation prior to the closure (RI_2014/VI_2011; *F_ST_* = 0.0006, P = 0.14) to statistically significant, although still very low, differentiation following the closure (RI_2023/VI_2020; *F_ST_* = 0.0021, P < 0.001). Prior to the closure, pairwise *F_ST_* values based on neutral loci ranged from 0.0005 (VI_2011/RI_2014) to 0.0976 (Figure 4). Following the closure, the minimum pairwise *F_ST_* increased slightly, with values ranging from 0.0021 to 0.0976, reflecting the emergence of low but significant differentiation between these two sites.

### Genetic diversity across time

Temporal changes in observed heterozygosity revealed contrasting trends across sites that did not follow a clear geographic pattern. Three sites (DO, GA and RI) showed a significant increase in observed heterozygosity (H_O_) following fishery closure, whereas the remaining two sites (VI and DE) exhibited a significant decline over the same period (Figure 5). These opposing trajectories suggest that the genetic consequences of fishery closure and reduced harvesting pressure were spatially heterogeneous and not driven by broad-scale regional processes. Complementary diversity metrics reinforced this site-specific pattern: VI was the only site showing a significant decrease in both expected heterozygosity (H_E_) and allelic richness (A_R_), while allelic richness increased significantly at RI. The remaining sites did not show significant changes in these additional metrics.

**Figure 5.**
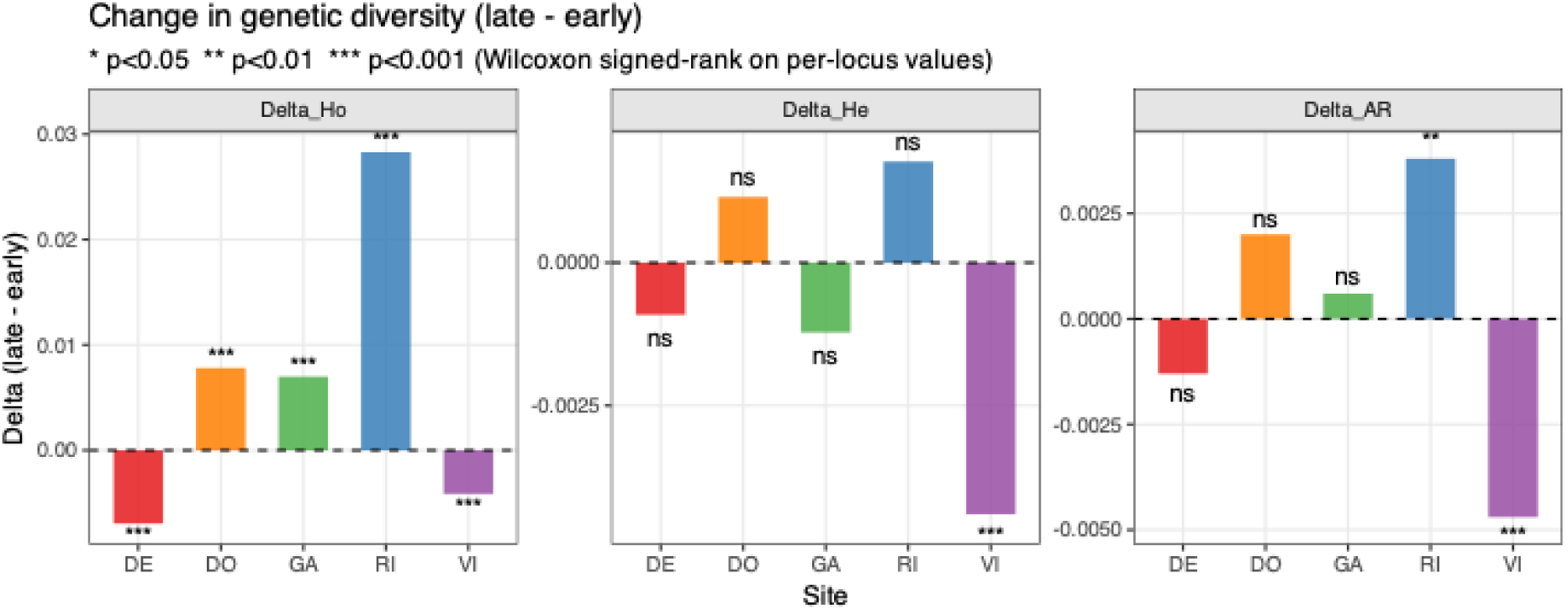
Genetic diversity changes over time. Observed heterozygosity, expected heterozygosity and allelic richness before and after fishery closure are shown for the five sampling locations with temporal replicates. Statistical tests were performed between temporal replicates, and corresponding P-values are indicated.

### Effective population size across regions and time

At the regional scale, estimated *Ne* increased in the Mediterranean Sea and declined in the Atlantic between the pre- and post-closure sampling periods. Mediterranean *Ne* rose from 2,861 (95% CI: 2,390-3,564) in 2014 to 4,225 (95% CI: 3,366-5,669), representing an approximately 1.5-fold increase. Conversely, Atlantic *Ne* declined from 1,174 (95% CI: 1,139-1,210) in the pre-closure period (2011-2014) to 718 (95% CI: 707-730) in the post-closure period. These contrasting temporal patterns should be interpreted cautiously, as they may reflect differences in recruitment dynamics, sampling scale, connectivity, local management history or estimator sensitivity rather than direct causal effects of fishery closure.

Using temporal methods, contemporary *Ne* estimates retained for the five reported locations were consistently low, ranging from 54.2 in GA to 97.7 in DO using the Pollak estimator (Table 3). Such low values are characteristic of species with high variance in reproductive success and should be interpreted with caution as point estimates. Notably, the highest retained Pollak estimates were recorded at DO and RI, whereas the lowest estimate was recorded at GA. These estimates should be interpreted as indicators of potential vulnerability rather than as direct evidence of demographic collapse or recovery.

**Table 3.** Location-specific temporal estimates of effective population size (*Ne*) for the five sampling locations with retained estimates. The number of loci analysed (n_loci), the number of individuals used in each sampling year (n1, n2) and the number of generations assumed (tgen) are shown. Pollak estimates are reported with confidence intervals; Nei/Tajima and Jorde/Ryman estimates are reported as point estimates.

| Site | Population_pair | tgen | n1 | n2 | n_loci | Pollak | Nei/Tajima | Jorde/Ryman |
| --- | --- | --- | --- | --- | --- | --- | --- | --- |
| DE | DE_2014/DE_2022 | 7 | 31 | 29 | 6274 | 70.6 (66.4 - 74.8) | 113.2 | 70.4 |
| DO | DO_2014/DO_2022 | 7 | 26 | 30 | 6584 | 97.7 (90.9 - 104.4) | 173.5 | 97.6 |
| GA | GA_2014/GA_2022 | 7 | 21 | 26 | 6337 | 54.2 (51.1 - 57.3) | 86.5 | 54 |
| RI | RI_2014/RI_2023 | 9 | 25 | 31 | 6700 | 93.3 (88 - 98.7) | 151.2 | 93 |
| VI | VI_2011/VI_2020 | 6 | 29 | 27 | 6620 | 73.5 (68.8 - 78.2) | 124.2 | 73.4 |

### Seascape genomics

At a broader spatial scale, the distance-based redundancy analysis (db-RDA) conducted on all samples demonstrated that spatial structure accounted for a greater proportion of genetic variance at neutral loci (R² adj = 4.80%, P-value < 0.001) than environmental variables (R² adj = 2.50%, P-value < 0.001). Several environmental predictor variables (PC1, PC2, PC3) exhibited strong correlations with spatial eigenvectors (MEM2, MEM10, MEM11), with all pairwise correlations exceeding 0.7, indicating substantial spatial structuring of the environmental signal. To address collinearity, variables were selected according to their explanatory power in univariate linear models, resulting in the retention of five variables in the final dbRDA model: PC1, PC3, MEM3, MEM4 and MEM9. All variance inflation factor (VIF) values were below 10, confirming an acceptable level of multicollinearity. Partial db-RDA further revealed that environmental factors alone were not statistically significant (P-value = 0.116), whereas spatial factors remained significant after removing the environmental component, still explaining 2.3% of the genetic variation (P-value < 0.001). This pattern suggests that the environmental signal is largely spatially structured and that geography captures genetic variance beyond what environmental variables alone can explain at neutral loci - consistent with the predominant role of isolation by distance and restricted gene flow in shaping neutral genetic structure across sampling locations.

The distance-based redundancy analysis (db-RDA) on the 649 candidate outlier SNPs revealed that spatial structure explained a greater proportion of genetic variation (R² adj = 19.0%, P-value < 0.001) than environmental factors alone (R² adj = 10.4%, P-value < 0.001). For the spatial component, MEM2, MEM3, MEM4, MEM9 and MEM11 were retained as significant predictors, while PC1, PC2 and PC3 were retained for the environmental component. Partial db-RDA further revealed that environmental factors lost statistical significance once spatial structure was accounted for, whereas spatial factors remained strongly significant after removing the environmental component (R² adj = 8.7%, P-value < 0.001). This suggests that the environmental signal is largely spatially structured - that is, environmental variation co-varies with geographic position - and that space captures genetic variation beyond what environment alone can explain. The final model was statistically significant, with an adjusted R² of 19.1% (Figure 6).

**Figure 6.**
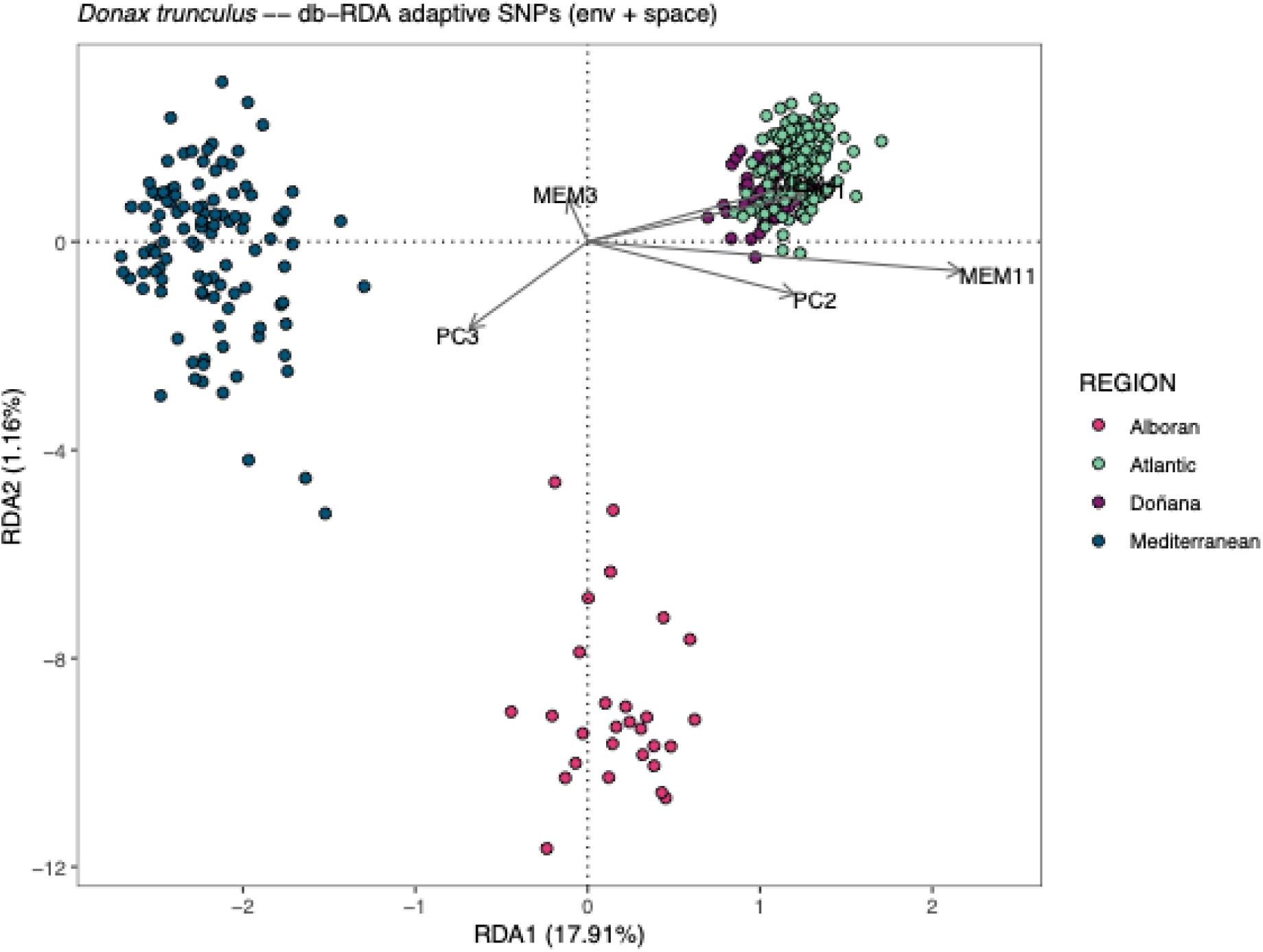
Genotype-by-environment analysis. Distance-based redundancy analysis showing the influence of spatial and environmental variables on candidate outlier genomic variation. The location of individuals in the ordination space reflects their relationship with explanatory variables based on multilocus genotypes at the 649 pcadapt candidate outlier loci.

Regarding the environmental predictors, PC1 - predominantly influenced by phosphate, dissolved inorganic carbon, and chlorophyll *a* - was associated with higher concentrations in Atlantic samples, reflecting the generally more productive and nutrient-rich conditions of the Atlantic region. In contrast, PC3 - characterised by human population density (PDNE), proportion of surrounding land cover (LAND), and a negative contribution of nitrate - showed elevated values in Mediterranean samples, potentially reflecting the stronger anthropogenic influence and distinct hydrographic conditions of that basin.

## Discussion

This study represents the first seascape genomics assessment of the commercially important wedge clam, *D. trunculus*, along the Iberian Peninsula coast, and the first to incorporate temporal replicates to evaluate the stability of genetic structure over time. Using genome-wide SNP data, we uncovered a temporally stable genetic structure marked by distinct spatial discontinuities. Although overall genetic differentiation was moderate to low (*F_ST_* < 0.1), our analyses revealed clear stock boundaries and fine-scale population structuring, confirming previous insights derived from microsatellite markers (Marie *et al*, 2016; Nantón *et al*, 2017).

### Temporal stability of genetic connectivity

Understanding the evolutionary consequences of anthropogenic pressures on natural populations is vital for assessing their evolutionary trajectories and predicting long-term resilience. In this study, our results suggest that gene flow and population connectivity remained largely stable between samples collected before and after fishery closure or reduced harvesting activity in most wedge clam fisheries in Spain. This pattern is consistent with limited detectable allele-frequency change over the time frame examined and agrees with Pinsky *et al* (2021), who reported temporal genomic stability in Atlantic cod despite decades of intensive exploitation, based on temporal replicates spanning over 70 years (1940-2013). The observed temporal genetic homogeneity may reflect the buffering effects of high reproductive output, planktonic larval dispersal and gene flow in marine invertebrates. A notable exception was the comparison between Ría de Arousa (RI) and Vilarrube (VI), where genetic differentiation was non-significant prior to the fishery closure but became statistically significant thereafter, suggesting possible recent changes in connectivity or localised demographic processes that warrant further investigation. However, given the subtle magnitude of this shift, these results would benefit from additional temporal sampling to confirm whether this represents a genuine biological trend.

### Isolation between regions

A biogeographic break between the Atlantic Ocean and the Mediterranean Sea has been documented across a wide range of species (Patarnello *et al*, 2007) and more recently confirmed through genomics approaches combined with large-scale sampling designs (Leiva *et al*, 2023). This break, located in the Alboran Sea, is associated with a semi-permanent dynamic front formed by the confluence of distinct water masses. The contrasting hydrographic and environmental conditions on either side of this front are thought to contribute to the observed genetic divergence. Prior to entering the Mediterranean through the Gibraltar Strait, Atlantic waters recirculate near Cape Trafalgar under the influence of strong tidal currents and an underwater ridge (Sala *et al*, 2023), creating a cold-water retention feature that may restrict larval dispersal and migration in some species, thereby acting as a barrier to gene flow (Sánchez-Garrido and Nadal, 2022). Our data corroborate this pattern, revealing significant genetic differentiation among individuals from the Atlantic Ocean, the Alboran Sea and the north- western Mediterranean coast, and confirming this biogeographic break as an important boundary for genetic connectivity in wedge clams.

Previous microsatellite-based studies identified three broad management units in *D. trunculus* - the Atlantic Ocean, the Balearic Sea, and the Alboran Sea - based primarily on neutral genetic differentiation (Marie *et al*, 2016; Nantón *et al*, 2017). Building on this framework, our analyses of candidate outlier SNP loci provide finer resolution by further differentiating populations within the Iberian Atlantic, particularly between southern Atlantic populations and those from the central/northern Atlantic. These results therefore refine, rather than replace, the management units proposed by earlier studies and highlight the importance of incorporating potentially adaptive genomic variation when delineating conservation and fisheries-management units for *D. trunculus*.

### Local adaptation

Seascape genomics seeks to identify potential environmental drivers of adaptive genetic variation by integrating genomic data with spatial environmental information across marine landscapes (Liggins *et al*, 2019; Riginos and Liggins, 2013). Applying this framework, our outlier-based db-RDA suggests that genomic variation at candidate loci is strongly spatially structured and partly associated with environmental gradients, including dissolved inorganic carbon, seafloor temperature and primary productivity (chlorophyll-a). However, because environmental variables covary with geographic distance and major oceanographic discontinuities, these results should not be interpreted as definitive evidence that these variables directly drive selection. Rather, they indicate candidate spatial-environmental associations that require further testing through additional sampling, experimental approaches or functional validation. Similar environmental associations have been reported in seascape-genomic studies of marine invertebrates, including links with temperature and chlorophyll concentration (Hollenbeck *et al*, 2022; Nielsen *et al*, 2020; Van Wyngaarden *et al*, 2018), and temperature-driven gene-expression differences between Atlantic and Mediterranean populations of another clam species, *Ruditapes decussatus* (Saavedra *et al*, 2023). At a broader taxonomic scale, comparative analyses across 195 bivalve species have also demonstrated significant influence of chlorophyll concentration and temperature on growth dynamics (Saulsbury *et al*, 2019). Together, these findings support the need to consider both spatial structure and possible environmental associations when defining conservation and management units for *D. trunculus*.

### Contrasting trends in genetic diversity over time

Temporal changes in genetic diversity were site-specific rather than regionally uniform. Observed heterozygosity increased at DO, GA and RI, but declined at VI and DE. However, complementary diversity metrics suggested that the strongest broader changes were concentrated at RI and VI: allelic richness increased significantly at RI, whereas VI showed decreases in both expected heterozygosity and allelic richness. These results indicate that changes in observed heterozygosity at DO, GA and DE should be interpreted cautiously, as they may reflect shifts in heterozygote frequency rather than broad gains or losses of genetic diversity. At RI, the increase in allelic richness is consistent with either local demographic improvement or increased immigration, whereas the decline detected at VI may indicate localised genetic erosion or recruitment variability. RI is an area with low fishing quotas and moderate fishing pressure, and immigration from nearby, less impacted sites may have contributed new individuals to this population. In contrast, Vilarrube is the last exploited wedge clam bank in Galicia. Despite this ongoing exploitation, wedge clam production has generally declined in Galicia in recent years, which has motivated research into restocking through hatchery rearing, using broodstock sourced precisely from the natural bank of Vilarrube. These contrasting patterns suggest that fishing pressure and site-specific management may shape trajectories of genetic diversity, with intensively and continuously exploited sites such as Vilarrube potentially more vulnerable to genetic erosion than sites subject to lighter, more diffuse harvesting such as RI. Yet, the absence of strong temporal changes in genetic diversity does not mean that demographic impacts are absent, because genomic erosion may become detectable only after a time lag, particularly in high-fecundity marine invertebrates with gene flow and large historical population sizes. Some depleted populations can retain high genetic diversity, sometimes exceeding levels observed in apparently healthy fisheries, suggesting that factors beyond exploitation intensity - including life history traits, recruitment variability and gene flow - can modulate demographic responses to harvesting pressure (Wooldridge *et al*, 2024). Continued monitoring and enforcement of conservation measures will therefore be important, particularly at sites where localised exploitation remains ongoing.

### Temporal shifts in effective population size

Overexploitation and fishery-induced demographic bottlenecks can reduce effective population size (*Ne*) in marine species, with cascading consequences for genetic diversity and long-term adaptive potential (Ciannelli *et al*, 2013; Pukk *et al*, 2013), although this pattern is not universal (Cuveliers *et al*, 2011). Here, temporal estimates of *Ne* showed contrasting regional patterns: estimated *Ne* increased in the Mediterranean Sea and declined in the Atlantic between the pre- and post-closure sampling periods. Nevertheless, these patterns should not be interpreted as direct evidence that fishery closure caused recovery or decline. Rather, they indicate temporal changes in estimated *Ne* that may reflect local recruitment dynamics, connectivity, sampling scale, or among-population differences in life history. First, differences in the intensity and continuity of fishing pressure between basins could differentially affect *Ne*, with sustained, spatially concentrated exploitation in the Atlantic - where VI now represents the last productive bank in Galicia - potentially accelerating genetic erosion, whereas the earlier closure of several Mediterranean fisheries (e.g., Murcia in 1998, the Balearic Islands in 2003, and the Valencian Community in 2017) may have allowed a partial relaxation of fishing-induced bottlenecks. Second, hatchery-based restocking efforts using broodstock sourced from a limited number of Atlantic donor banks could further reduce *Ne* through a Ryman-Laikre effect (Ryman and Laikre, 1991), independent of natural demographic trends. Third, differences in metapopulation connectivity between basins may influence estimated *Ne*, as gene flow from less-exploited source populations can inflate local *Ne* estimates in ways that do not necessarily reflect true increases in local effective population size.

Fine-scale temporal *Ne* estimates in *D. trunculus* for the five locations for which temporal estimates were retained were low, generally falling near or below 100 individuals. Although such values warrant management attention, they should be interpreted cautiously because *Ne* estimates can be sensitive to sample size (here sample size < 50), spatial scale, overlapping generations, variance in reproductive success and assumptions about generation time. These estimates are therefore best viewed as indicators of potential vulnerability requiring further monitoring, rather than as definitive evidence of imminent extinction risk. Conservation guidelines suggest that low *Ne* can increase the risk of inbreeding and reduce long-term adaptive potential (Frankham *et al*, 2014; Frankham *et al*, 2010), but applying these thresholds to high-fecundity marine invertebrates requires care. In *D. trunculus*, among-population variation in generation time may also influence the translation of *Ne* estimates into ecological and demographic terms.

### Translocation plans

Conservation translocations - defined as deliberate movement and release of organisms beyond their historical or current natural range with the aim of establishing or reinforcing populations of species of concern (Griffith *et al*, 1989) - remain a topic of considerable debate (Müller and Eriksson, 2013; Ricciardi and Simberloff, 2009). The relatively uniform genetic structure and temporal stability observed across northern and central Atlantic populations - Vilarrube (VI), Ría de Arousa (RI), and Setúbal (SE) - suggest that, from a genetic perspective, assisted movements among these locations may carry lower risk than transfers among highly differentiated regions. Any such intervention should be considered only after a comprehensive management assessment demonstrates that demographic reinforcement is necessary and that local recovery cannot be achieved through reduced fishing pressure, habitat protection or improved enforcement alone. Any pilot reinforcement should also be conditional on pathogen and parasite screening, habitat-suitability assessment, evaluation of donor-population status to avoid compromising source populations, and pre- and post-release genetic monitoring. Conversely, the genetic distinctiveness of the Alboran Sea and north-western Mediterranean/Balearic Sea clusters, as well as the southern Atlantic population from Doñana, argues against cross-regional translocations except under exceptional circumstances and after rigorous risk assessment. Transfers across these differentiated units could disrupt locally adapted gene complexes, increase the risk of outbreeding depression and compromise the evolutionary integrity of recipient populations (Frankham *et al*, 2011). The principal implication of our results is not that translocation should be pursued, but that genomic data can provide a principled framework for defining the boundaries within which any future reinforcement actions, if deemed necessary, should be carefully evaluated and monitored.

## Conclusions

Our temporal seascape-genomic analysis of *D. trunculus* reveals a largely stable genomic structure across the Iberian Peninsula, structured primarily by persistent biogeographic discontinuities between the Atlantic Ocean, Alboran Sea and north-western Mediterranean/Balearic Sea, while also identifying finer-scale Atlantic differentiation and site-specific temporal changes in genetic diversity. These results refine previous microsatellite-based management units and show that neutral SNPs, candidate outlier loci, temporal diversity metrics and *Ne* estimates provide complementary but distinct information for conservation and fishery management. More broadly, this study shows that temporal seascape genomics can separate persistent biogeographic structure from short-term, site-specific genomic change in exploited high-fecundity marine invertebrates. This framework is therefore useful not only for delineating management and potential reinforcement units in *D. trunculus*, but also for assessing how marine resources may be impacted at the genomic level by exploitation, closure and spatially heterogeneous recovery.

## Acknowledgments

We are grateful to Conselleria d’Agricultura, Aigua, Ramaderia i Pesca de la Generalitat Valenciana, and Consejería de Agricultura, Pesca, Agua y Desarrollo Rural (Junta de Andalucía) for facilitating sample collection.

## Author contributions

CR, LB, MH and MB designed the study. CR and MH wrote the project proposal and obtained funding. MB, CS, MR, SF, and MD contributed to the preparation of the project proposal and the execution of the work. MB, CS, MR, SF, MD, LS and AI coordinated the collection of samples. LB performed the bioinformatic analyses and prepared the figures. LB and CR wrote the first draft of the manuscript. All authors contributed to the final version and approved the submitted manuscript.

## Data availability statement

The dataset has been deposited in Zenodo: https://doi.org/10.5281/zenodo.21396105.

---

**Figure S1.**
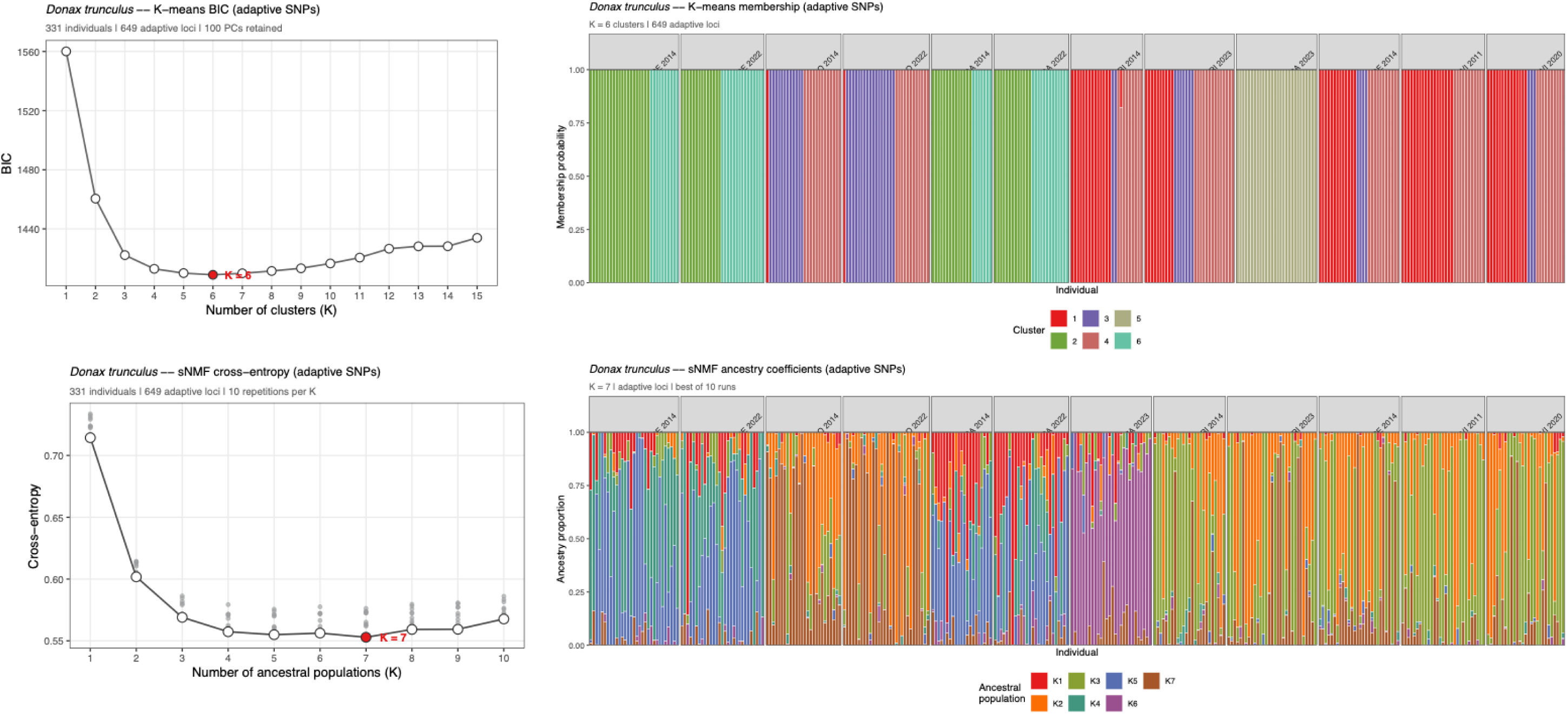

**Table S1.**
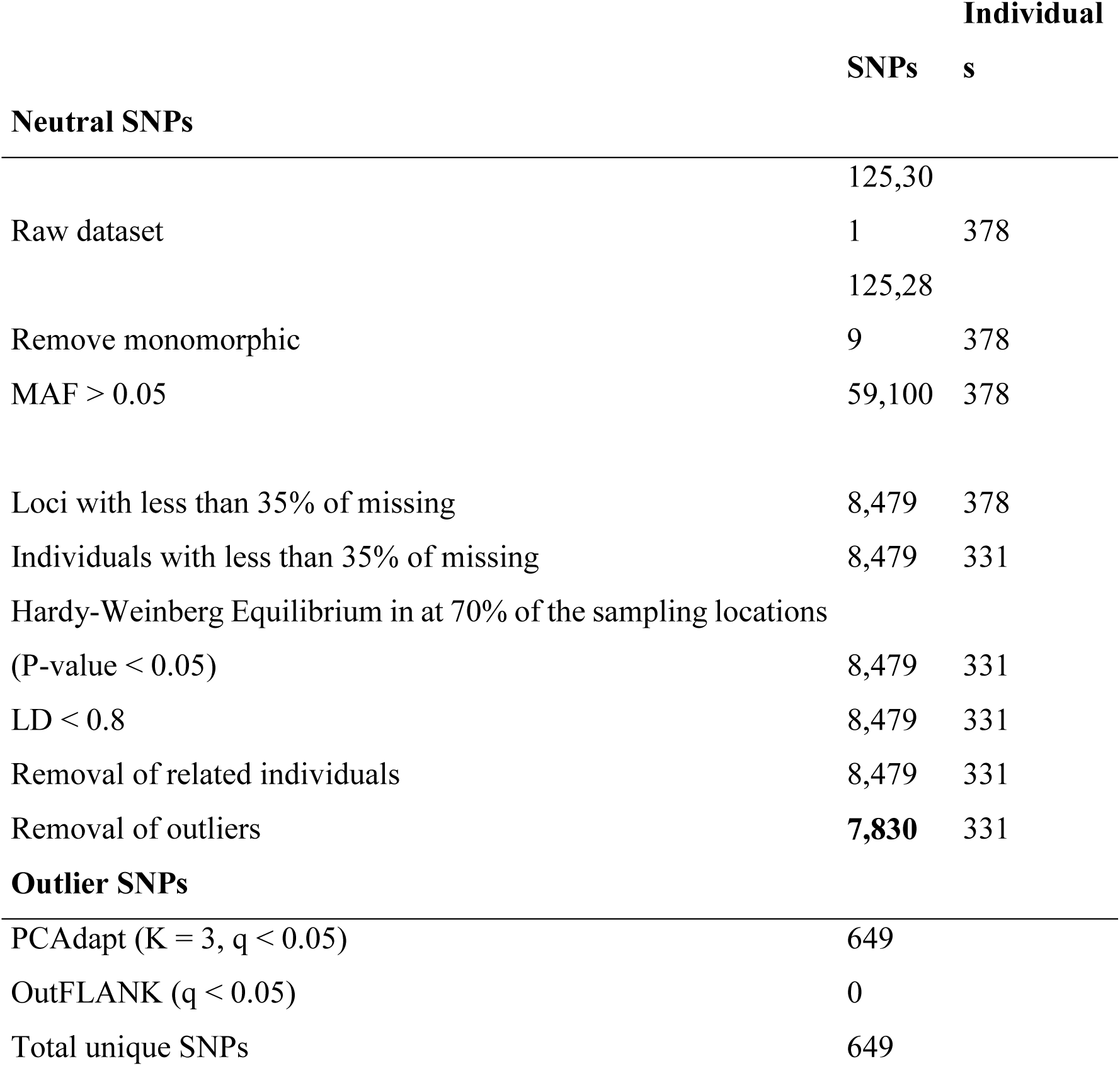
Filtering steps undertaken with subsequent individuals and SNPs remaining per filtering stage.

